# DriverPower: Combined burden and functional impact tests for cancer driver discovery

**DOI:** 10.1101/215244

**Authors:** Shimin Shuai, Steven Gallinger, Lincoln Stein, on behalf of the PCAWG Drivers and Functional Interpretation Group and the ICGC/TCGA Pan-Cancer Analysis of Whole Genomes Network

## Abstract

We describe DriverPower, a software package that uses mutational burden and functional impact evidence to identify cancer driver mutations in coding and non-coding sites within cancer whole genomes. Using a total of 1,373 genomic features derived from public sources, DriverPower’s background mutation model explains up to 93% of the regional variance in the mutation rate across a variety of tumour types. By incorporating functional impact scores, we are able to further increase the accuracy of driver discovery. Testing across a collection of 2,583 cancer genomes from the Pan-Cancer Analysis of Whole Genomes (PCAWG) project, DriverPower identifies 217 coding and 95 non-coding driver candidates. Comparing to six published methods used by the PCAWG Drivers and Functional Interpretation Group, DriverPower has the highest F1-score for both coding and non-coding driver discovery. This demonstrates that DriverPower is an effective framework for computational driver discovery.

## Introduction

Cancer drivers are somatic genetic alterations that confer selective advantages to tumour cells^1,2^. Identification of cancer drivers is a crucial yet challenging task in cancer genomics research^3,4^. There are multiple challenges. First, driver mutations generally account for only a small fraction of the somatic variations found in a typical tumour, the rest being innocent bystander “passenger” mutations^5^. Second, there is substantial intra- and inter-tumoural heterogeneity in most cancers^6^. Both across different tumour types and across different genomic regions within the same tumour, the background mutation rate can vary over several orders of magnitude.

The advent of large scale cancer whole-genome sequencing (WGS) data, such as the ˜2,600 tumour and matched normal whole genomes from the ICGC/TCGA Pan-Cancer Analysis of Whole Genomes (PCAWG) project, has made it possible to explore the role of driver events in non-coding regions. However, identifying non-coding driver events in WGS creates new challenges. First, while the functional impact of somatic mutations in the coding regions of genes is fairly straightforward to predict, much less is known about the effect of mutations on non-coding regions of the genome. Second, only ˜1% of somatic mutations detected in PCAWG WGS data are exonic, adding substantially more mutations and regions to be tested and demanding more careful control of type I and type II errors than whole exome sequencing. At present, only a limited number of non-coding drivers are known, the primary examples being the *TERT* promoter for multiple tumour types and the *TAL1* enhancer for T cell acute lymphoblastic leukaemia^7,8^.

Most state-of-the-art methods identify drivers by detecting signals of positive selection either through mutational burden tests, which compare the rate of mutations observed in a region of the genome to what is expected from the background mutation rate, or functional impact tests, which identify putative driver mutations based on a higher-than-expected rate of changes that are predicted to alter the function of genomic elements^3,6^. Mutational burden tests work best for calling frequently recurrent driver events and perform poorly when applied to rare driver events. In contrast, functional impact tests fail to find drivers in genomic elements that are poorly understood or annotated. To maximize accuracy, we combined the two mutation significance testing methods to develop DriverPower (**Fig. 1a**), a framework for identification of coding and non-coding cancer drivers using mutational burden and functional impact scores.

**Figure 1.**
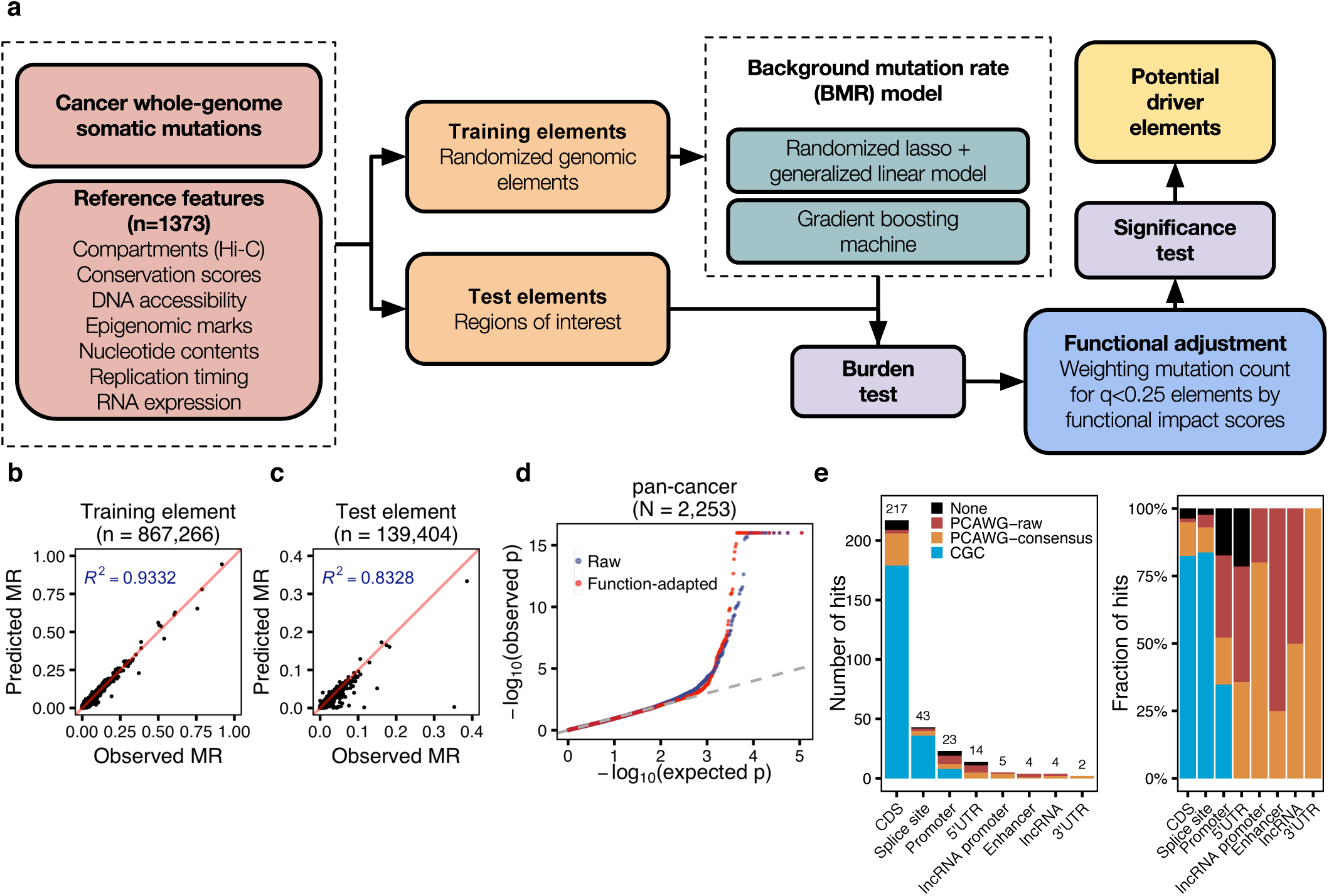
Summary of method and results. (**a**) DriverPower overview. (**b**,**c**) For the training and test element sets, comparison of the predicted (Y-axis) and observed (X-axis) mutation rate in the pan-cancer cohort. (**d**) The raw and function-adapted p-value quantile-quantile (QQ)-plot for all test elements in the pan-cancer cohort. Function-adapted p-values are p-values with the incorporation of functional impact scores. (**e**) Number and fraction of non-coding driver candidates called by DriverPower contained within three reference driver sets (CGC, PCAWG-consensus or PCAWG-raw). For each element type, the number of candidates is also shown above the bar.

## Results

To evaluate DriverPower, we took WGS somatic variant data derived from 2,583 high-quality donors from the PCAWG project^9^. After removing hypermutated samples, 2,514 donors with 24,715,214 somatic single nucleotide variants (SNV) and small indels were used for driver element identification. We analysed this data both as a single pan-cancer data set, as well as a series of 29 tumour type-specific cohorts (**Supplementary Table 1**).

### Features predictive of background mutation rate

Among all tumour cohorts, we observed substantial variability in the observed mutation rate at the tissue, donor and locus levels (**Supplementary Fig. 1** and **2**). Accurate driver detection requires an accurate estimate of background mutation rate across the tumour genome, taking into account the extensive variability among tumour types, donors and genomic regions. DriverPower tackles this issue by modelling the background mutation rate (BMR) using numerous genomic features that co-vary with the localized background mutation rate. We collected 1,373 features from three public data portals (**Supplementary Table 2**): the ROADMAP Epigenomics project, the ENCODE project and the UCSC genome browser^10,11,12^. These features covered seven main categories: conservation, DNA accessibility, epigenomic marks, nucleotide contents, replication timing, RNA expression and genome compartments. As expected, we found extensive multicollinearity among features. Most features (1368/1373) are significantly correlated with pan-cancer genome-wide mutation rates (Spearman’s rho test q*<*0.1; **Supplementary Fig. 3**).

### Background mutation rate model

We investigated two algorithms for modelling the BMR based on genomic features. The first algorithm was randomized lasso followed by binomial generalized linear model (GLM). The alternative algorithm was the gradient boosting machine (GBM), which is a non-linear and non-parametric tree ensemble algorithm^13^. To evaluate both BMR modelling algorithms, we made non-overlapped 1 megabase pair (Mbp) autosomal elements (n=2,521) as well as training genomic elements (n=867,266) by sampling genomic coordinates randomly. The number of mutations per element was then predicted with 5-fold cross-validation.

When evaluated using 1Mbp autosomal elements, we found that both algorithms could accurately predict the background mutation rate (**Supplementary Fig. 4** and **5**). In high mutational burden tumour cohorts, we observed essentially no difference between two algorithms, however GBM consistently outperformed GLM when applied to low mutational burden tumour cohorts (**Supplementary Fig. 6**). When evaluated on the training element set, in which the size of element varies from 100 bp to 1 Mbp, the prediction accuracy drops due to higher BMR variability, especially for low mutational burden tumour cohorts such as Myeloid-MPN and CNS-PiloAstro (**Supplementary Fig. 6**). However, for large cohort such as the pan-cancer set (N=2,253), around 93% of the mutation rate variance on the training set is explained by either model (**Fig. 1b**). The model still shows excellent performance when applied to the test element set, explaining 83% of the mutation rate variance on the pan-cancer cohort (**Fig. 1c**).

Both the randomized lasso algorithm and the GBM can be used to rank feature importance in different ways. Feature selection ranking from both methods confirmed that H3K9me3 (associated with heterochromatin), replication timing and H3K27ac (or its antagonistic histone mark H3K27me3) are the most important groups of predictors for BMR (**Supplementary Fig. 7** and **Supplementary Table 2**)^14^. Consistent with previous results, we found that features from tumour cell lines with similar cell-of-origin to the primary tumour type are frequently selected^15^. For example, replication timing from liver cancer cell line HepG2 was selected as a feature for the BMR in hepatocellular carcinoma (Liver-HCC), while replication timing in MCF7 (breast cancer) and SK-N-SH (neuroblastoma) were selected for breast adenocarcinoma (Breast-AdenoCA) and glioblastoma (CNS-GBM), respectively (**Supplementary Fig. 8**).

### Functional adjustment

In most burden-based methods mutations are equally weighted. However, not all mutations have the same functional consequences. To incorporate functional consequence information, DriverPower implements a posterior functional adjustment. The functional adjustment step upweights mutations with high predicted functional impact. While the DriverPower framework can potentially work with any functional scoring scheme, in the current implementation we measured the functional impact using four published scoring schemes: the CADD^16^, DANN^17^, EIGEN^18^ and LINSIGHT^19^ scores. Although different training data, assumptions and algorithms are used by different scores, we found those scores to be consistent at the element level (**Supplementary Fig. 9**). We used the average weight of all four scores in the remainder of the manuscript unless otherwise specified.

### Candidate driver event discovery

To evaluate the DriverPower algorithm, we first employed three simulated variant sets generated by the PCAWG Drivers and Functional Interpretation Group (PDFIG) to examine type I and type II errors. We expected to identify no drivers as all three simulated datasets are reshuffles of observed mutations. In general, we observed no inflation or deflation in simulations and only 8 significant hits were identified in ˜11M statistical tests (**Supplementary Fig. 10**). We then used the observed PCAWG dataset to discover drivers within multiple coding and non-coding element sets identified by the PDFIG, spanning 3.7% (˜113 Mbp) of the human genome.

We benchmarked our results against reference driver element sets and driver candidates called by six other published methods. Among the six methods, ExInAtor^20^, ncdDetect^21^ and LARVA^22^ use only mutational burden information; oncodriveFML^23^ uses only functional biases; while MutSig^24^ and ActiveDriverWGS^25^ model both mutational burden and functional consequence but not through functional impact scores. Three reference driver element sets were used: the COSMIC Cancer Gene Census (CGC)^26,27^, the PCAWG raw integrated driver candidates (PCAWG-raw) and the PCAWG consensus driver candidates (PCAWG-consensus). The CGC is a catalogue of driver genes for which mutations have been causally implicated in cancer and was used as the gold standard set (*i.e.*, used in the calculation of precision and recall) for coding and splice site drivers. PCAWG-raw is an integration of driver elements called by 12 different driver detection methods on the same data we used here. PCAWG-consensus is a conservative set derived from the PCAWG-raw by applying multiple stringent filters to control the false discovery rate; in particular, the majority of non-coding candidates from lymphoid tumours and skin melanomas is excluded from this set because of hyper-mutational processes in these tumour types that create prominent mutational hotspots^28,29,30^. For the same reason our analysis of non-coding regions for tumour-specific and the pan-cancer cohorts excluded melanoma and lymphoma.

Overall, we observed well calibrated p-values in DriverPower’s results with or without functional adjustment (**Fig. 1d** and **Supplementary Fig. 10**) and a high accuracy for both coding and non-coding driver discovery (**Fig. 1e**, **Supplementary Table 3**). For protein coding regions (CDS), DriverPower found 217 significant driver candidates. Since a gene (e.g., *TP53*) can be driver in multiple cohorts, the unique number of genes was 131. The precision of the algorithm’s driver calls was high. Among the driver genes called by DriverPower, 82.5% (179/217) of all genes were present within the CGC. For non-CGC genes, 27 and 3 genes were present within PCAWG-consensus and PCAWG-raw, respectively. Thus, only 3.7% (8/217) coding driver candidates called by DriverPower were not contained within any reference gene sets. As expected, incorporation of functional information increased both precision and recall in coding driver discovery (**Fig. 2a** and **Supplementary Fig. 11**). For example, in pancreatic ductal adenocarcinoma (PancAdenoCA; N=232), the addition of functional adjustment to the algorithm resulted in a gain of three additional drivers (*ACVR1B*, *RBM10* and *ZFP36L2*) and the loss of one likely false positive genes (*FAU*) (**Fig. 2a**). Without the use of functional information, the overall precision dropped to 74.6% (156/209) for CGC genes and 88.5% (185/209) for CGC/PCAWG genes. When compared to six other methods using the same 26 non-melanoma/lymphoma cohorts and CGC as the gold standard set, DriverPower (precision=0.84; recall=0.79) had the highest F1score (0.81) (**Fig. 2b, c**). In our benchmark, sensitivity was a bottleneck for most methods (4/7 with recall*<*0.5). When compared to the method with highest recall, the widely used coding driver caller MutSig (precision=0.80; recall=0.80), DriverPower identified an additional 21 genes present in CGC (23 for MutSig; **Supplementary Fig. 12**).

**Figure 2.**
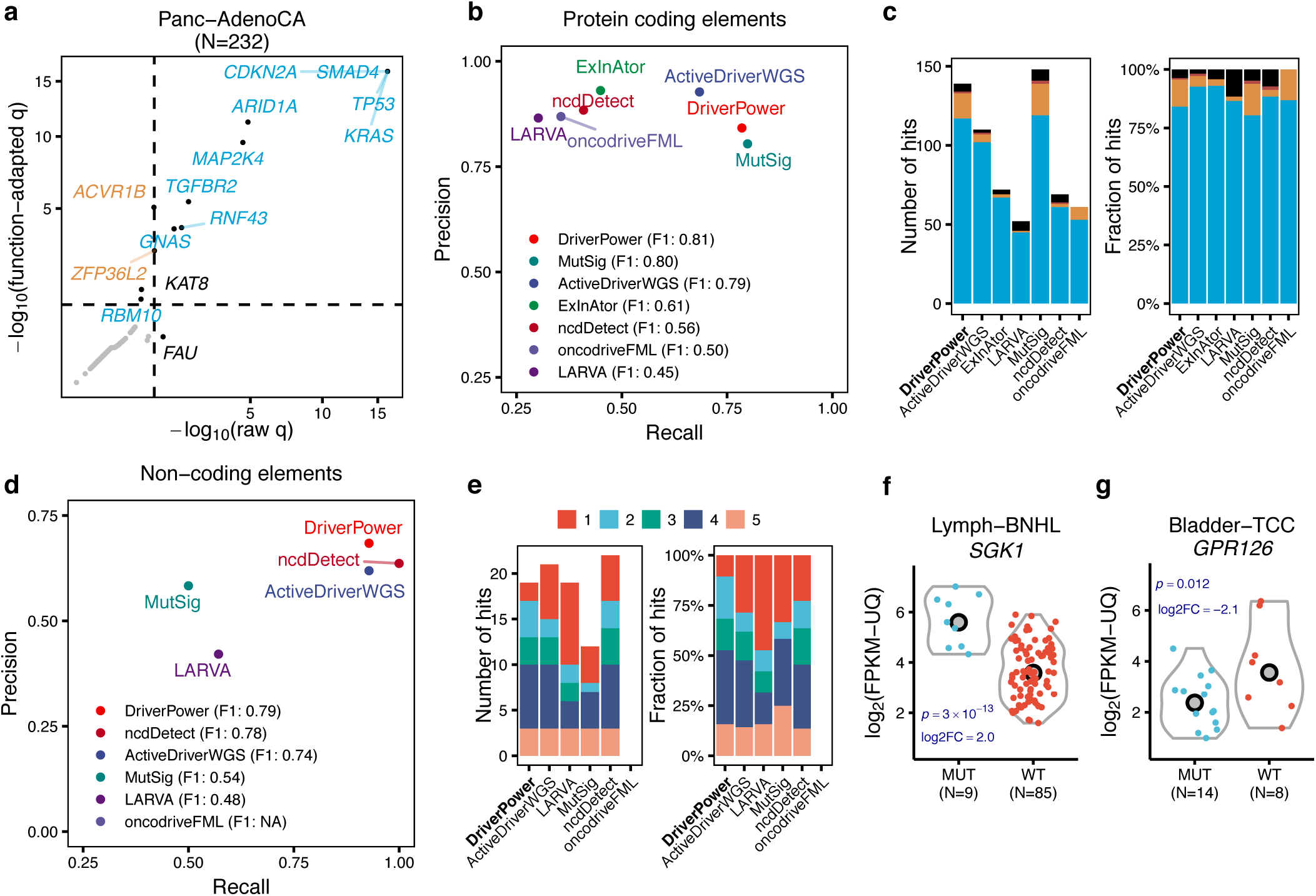
Benchmarking DriverPower driver discovery performance. (**a**) Comparison of CDS results with or without functional adjustment for Panc-AdenoCA. Dashed lines in (a) represent the q-value = 0.1 threshold. Function-adapted q-values are q-values with the incorporation of functional impact scores. Only significant genes are labelled (color legend is the same as **Fig. 1e**). (**b**,**c**) Benchmark results for coding genes compared to six other driver discovery methods. (**d**,**e**) Benchmark results for 3’UTR, 5’UTR, promoter and enhancer sets. (b) and (d) show the precision and recall for each method according to results of 26 tumour cohorts (no melanoma and lymphoma). (c) shows the number and fraction of coding driver candidates called by each method that are contained within reference gene sets. The colored columns in (c) correspond to different reference driver sets (color legend is the same as **Fig. 1e**). (e) shows the number and fraction of non-coding driver candidates called by each method that are also called by others. The colored columns in (e) correspond to the number of methods that agree on a driver candidate. (**f**) Differential expression analysis for the CDS and splice site of *SGK1* in Lymph-BNHL. (**g**) Differential expression analysis for the *GPR126* enhancer in Bladder-TCC. MUT indicates samples with mutated element and WT indicates samples without mutated element. Copy number corrected p-values from the likelihood ratio test and the log2 fold changes (log2FC) are shown in blue.

We next benchmarked DriverPower’s accuracy for non-coding driver events. For the prediction of driver events affecting the splice sites of coding genes, DriverPower called 47 significant candidates with 85.1% (40/47) within CGC. DriverPower (F1=0.91) also outperformed two recently published methods, ncdDetect (F1=0.65) and oncoDriverFML (F1=0.32), for splice site driver detection (**Supplementary Fig. 13**).

For the prediction of non-coding driver events in 3’UTRs, 5’UTRs, promoters and enhancers, DriverPower identified 19 candidates in non-melanoma/lymphoma tumour cohorts and 24 candidates in the pan-cancer cohort. Benchmarking results showed that DriverPower has the highest F1-score (0.79) among the six methods evaluated (**Fig. 2d, e**). Promoter and 5’UTR driver candidates called by DriverPower were associated with a total of 17 unique genes. Of these, one gene (*TERT*) was in CGC, four genes (*WDR74*, *HES1*, *MTG2* and *PTDSS1*) were in PCAWG-consensus, and six other genes were in PCAWG-raw. DriverPower also called two 3’UTR driver candidates in total, including *TOB1* in pan-cancer and *ALB* in Liver-HCC. Both candidates were present in the PCAWG-consensus. For enhancer regions, DriverPower identified two candidates: chr6:142,705,600-142,706,400 (linked to *GPR126*), and chr7:86,865,600-86,866,400 (linked to *TP53TG1*). Both enhancer elements were identified by PCAWG-raw; the *TP53TG1* enhancer was the only enhancer for non-melanoma/lymphoma tumours in the PCAWG-consensus set.

For long non-coding RNA (lncRNA) genes and their promoters, DriverPower found 9 candidates in total. Among them, 6 and 3 were contained within PCAWG-consensus and PCAWG-raw, respectively. These candidates targeted three unique lncRNAs: *RN7SK, RMRP* and *RPPH1*. The promoter of *RMRP* was significantly mutated in four cohorts (Breast-AdenoCA, Liver-HCC, Stomach-AdenoCA and pan-cancer) and has been nominated as a novel non-coding driver.

### DriverPower-exclusive driver candidates overview

A total of 11 coding and 17 unique non-coding candidates were exclusively identified by DriverPower (not present in either CGC or PCAWG-consensus; **Supplementary Table 4**). We sought to evaluate these exclusive driver candidates using literature evidence and correlative orthogonal data such as the effect of the variant on RNA-seq expression levels and the presence of somatic copy number alterations (SCNAs) and somatic structural variations (SVs) covering the same regions. On this basis, we found that many of the DriverPower-exclusive candidates are plausible cancer drivers.

Among protein coding genes, DriverPower identified *EEF1A2* (eukaryotic translation elongation factor 1 alpha 2) in the esophageal adenocarcinoma cohort (Eso-AdenoCA; 7/95 samples). All seven observed mutations were missense (**Supplementary Fig. 14a**). Although no RNA-seq data is available for Eso-AdenoCA samples, SCNA analysis indicated that *EEF1A2* is amplified in 69.5% (66/95) of Eso-AdenoCA samples (vs. 27.9% of non-Eso-AdenoCA samples; **Supplementary Fig. 14b**), suggesting a potential gain-of-function role in this cancer type. The amplification of *EEF1A2* (20q13.33) was also confirmed by the GISTIC2.0 (q=0.0006). The same 1Mbp locus detected by GISTIC2.0 was also amplified recurrently in other tumour types, including 73.1% of colorectal adenocarcinoma, 64.7% of stomach adenocarcinoma and 55.4% of ovarian adenocarcinoma. Supporting this hypothesis, previous studies have also demonstrated that *EEF1A2* is a putative oncogene in ovarian cancer and overexpressed in various tumour types^31,32,33^.

Another protein coding gene exclusively identified by DriverPower was *MEF2B* in B-cell nonhodgkin’s leukaemia (Lymph-BNHL; 8/105 samples). Among 9 observed mutations, 8 mutations were missense and one was a frameshift deletion (**Supplementary Fig. 14c**). RNA-seq data confirmed that mutated samples overexpressed *MEF2B* (copy number corrected p=0.011; **Supplementary Fig. 14d**). *MEF2B* (Myocyte enhancer factor 2B) has been identified in multiple WES studies^34,35,36^, and a previous study has also shown that *MEF2B* mutations can dysregulate cell migration in non-Hodgkin lymphoma^37^.

One splice site candidate exclusively called by DriverPower is *SGK1* (serum/glucocorticoid regulated kinase 1) in Lymph-BNHL. The same gene was also significant in DriverPower’s CDS result for Lymph-BNHL (**Supplementary Fig. 14e**), resulting in a total of 13.3% (14/105) Lymph-BNHL samples being affected by non-synonymous or splice site mutations in *SGK1*. *SGK1* is present in PCAWG-raw but was filtered out due to the large number of AID-related variants in this tumour cohort. However, differential expression analysis indicated that *SGK1* is significantly overexpressed in mutated Lymph-BNHL samples relative to non-mutated samples (copy number corrected p=3e-13; **Fig. 2f**). *SGK1* encodes a serine/threonine protein kinase that plays an important role in cellular stress response and its CDS has been nominated as a driver in earlier WES studies^35,36^. Another study has also demonstrated that the administration of an *SGK1* inhibitor induces apoptosis in lymphoma cell lines^38^. Together these data support a potential driver role for *SGK1* in Lymph-BNHL.

The *GPR126* (adhesion G protein-coupled receptor G6) enhancer candidate was filtered out from the PCAWG-raw set because of mutations in palindrome loops, which makes it unclear whether mutations in the *GPR126* enhancer are caused by mutational mechanism associated with palindrome loops or positive selection. We found that the *GPR126* enhancer is recurrently mutated in transitional cell carcinoma of the bladder (Bladder-TCC; 14/23 samples) and breast adenocarcinoma (Breast-AdenoCA; 8/195) (**Supplementary Fig. 14f**). *GPR126* is among the MammaPrint^®^ 70 gene panel used to predict the risk of breast cancer metastasis^39,40^. A study also shows that knockdown of *GPR126* can inhibit the hypoxia-induced angiogenesis in model organisms^41^. Differential expression analysis demonstrated that the *GPR126* is significantly downregulated in Bladder-TCC samples with enhancer mutations (copy number corrected p=0.012; **Fig. 2g**) relative to those carrying the wild type enhancer, suggesting a functional role for these mutations.

Several somatically altered histone genes have been implicated in human cancer, such as *H3F3A* (identified as a pan-cancer driver in this study), *H3F3B* and *HIST1H3B^42,43,44^*. DriverPower identified four histone genes as driver candidates in the pan-cancer cohort, two of which were absent from CGC or PCAWG-consensus: the 5’UTR of *HIST1H2AC* and *HIST1H2BD*. Previous studies have shown that the protein levels of the replication-dependent histone H2A variant *HIST1H2AC* (encoding histone H2A type 1-C) is decreased in chronic lymphocytic leukaemia patients and bladder cancer cells^45,46^, and the siRNA knockdown of *HIST1H2AC* increases cell proliferation and promote oncogenesis^46^.

Several other driver candidates exclusively called by DriverPower are associated with genes that may have a role in cancer. The highly expressed liver-specific gene *ALB* (albumin) is significant for somatic mutations affecting its CDS, splice site, 3’UTR and promoter in Liver-HCC; the splice site and promoter (under CADD scores) were discovered by DriverPower exclusively. Correlative evidence from DNA promoter methylation, gene expression and copy number alterations suggested that loss-of-function mutations in *ALB* are subject to positive selection in Liver-HCC as described in ref. xx (Driver group main paper). The CDS of *KAT8* (lysine acetyltransferase 8) was called by DriverPower in Panc-AdenoCA with 100% (5/5) missense mutations. As a histone modifier, *KAT8* has been shown to physically interact with *MLL1* and regulate known cancer drivers *ATM* and *TP53^47,48,49,50^*. Previous studies have also shown that *KAT8* is downregulated in gastric cancer^51^ and *KAT8* can suppress tumour progression by inhibiting epithelialto-mesenchymal transition^52^. The 5’UTR and promoter of *SRSF9* (serine and arginine rich splicing factor 9) was significant in DriverPower’s results for pan-cancer and not present in any reference driver sets. The protein encoded by *SRSF9* is part of the spliceosome; a previous study indicates that the proto-oncogene *SRSF9* is overexpressed in multiple tumours and that this overexpressi-on can cause the accumulation of β-Catenin^53^. The same study also showed that the depletion of *SRSF9* proteins could inhibit colon cancer cell proliferation.

In summary, 4/11 coding and 4/17 unique non-coding driver candidates exclusively called by DriverPower had some form of support from the literature or orthogonal evidence. If we assume that all the exclusive candidates that lack such evidence are false positives, then this puts an estimate of DriverPower’s false discovery rate across the PCAWG data set at 3.2% (7/217) for coding and 16.8% (16/95) for non-coding regions. However, this assumption is probably invalid as most of these lack-of-evidence candidates are also identified by other methods and present in PCAWG-raw. We acknowledge that lack-of-evidence candidates may contain false positive calls, but they may also contain previously unknown drivers. For example, the 5’UTR of *TBC1D12* in Breast-AdenoCA, which has been filtered out from the PCAWG-raw due to possible hypermutability, is called by all but one driver discovery methods and is reported as a putative cancer driver in previous studies because of two recurrent mutations in the Kozak consensus sequence involving in the initiation of protein translation^23,54^. Moreover, according to another recent study, the same *TBC1D12* candidate is still statistically significant in breast cancer even after removing hypermutations, but whether these mutations can alter protein translation in cancer is still undetermined^24^. Some lack-of-evidence candidates may also fit the mini-driver model of cancer evolution^55^. Unlike classical drivers, mini-drivers can only weakly promote and are not essential for tumour progression, hence present at a lower frequency in cancer cohorts. Further investigation is required to determine the role of lack-of-evidence candidates in cancer.

### DriverPower applied to whole exome sequencing

To demonstrate the robustness of DriverPower, we applied DriverPower to two public whole-exome sequencing (WES) datasets (**Supplementary Fig. 15**). Both WES datasets are processed differently than the PCAWG data and contain samples not included in the PCAWG study. For liver cancer, using models trained for Liver-HCC (N=314), DriverPower identified 14 coding drivers from 364 TCGA-LIHC samples (53 shared with Liver-HCC). All but one driver candidates were present within the CGC or PCAWG-consensus. For pancreatic adenocarcinoma, using models trained for Panc-AdenoCA (N=232), DriverPower identified 6 coding drivers from 180 TCGA-PAAD samples (no shared samples with the PCAWG study) and all corresponded to known driver genes.

## Discussion

Computational driver discovery is essential to distinguish driver from passenger mutations in the coding and non-coding regions of whole cancer genomes. Here we report DriverPower, a new framework for accurately identifying both types of driver mutation by combining mutational burden and functional impact information. The method takes advantage of the large somatic mutation sets produced by WGS technology to build an accurate global BMR model from more than a thousand genomic features. This contrasts with methods that build a local BMR model using selected or flanking regions. One advantage of this is that the method is not biased towards coding regions, but uses the same model for coding and non-coding cancer driver discovery. Another advantage is the method’s high degree of modularity. DriverPower can potentially work with any types of genomic element (contiguous or disjoint, coding or non-coding, proximate or distal to genes), any regression algorithms for modelling BMR and any functional impact score scheme. Although DriverPower is designed for WGS projects, it performs robustly in whole-exome sequencing strategies as well.

In comparison to the other driver discovery methods evaluated by the PCAWG Drivers and Functional Interpretation Group, DriverPower provides the best balance of precision and recall, although is not always the top ranked method when either metric is considered independently (**Fig. 2b**,**d**). As discussed in **Supplementary Note 1**, DriverPower is parameterized to allow for adjustment of the precision-recall trade-off; in this study, we selected conservative parameters that prioritize precision over recall especially for non-coding regions.

There are several ways in which the accuracy of DriverPower could be improved. One approach to improve recall is to take into the account the potential presence of negative (purifying) selection in the functional regions selected for testing. When the BMR model is trained, we use random genomic elements that are predominantly under neutral selection. However, the functional elements selected for testing are more likely to be under positive and/or negative selection^56^. The observed mutation rate reflects the balance between positive and negative selection, and negative selection at one site in the element will diminish the signal of positive selection at other sites, reducing the sensitivity of the method as a whole. To our knowledge, no driver discovery tool currently models the effect of negatively selected sites; future work aims to take this mechanism into account.

The precision of the method can also be improved. False positive driver calls may be caused by technical errors such as variant-calling artefacts that artificially increase the local mutation rate, or by biological processes that are not captured by the BMR model such as regional differences in activation-induced cytidine deaminase (AID) activity. These can potentially be mitigated by incorporating into the BMR model additional features relevant to the technical and biological processes. For example, incorporating read-level coverage, mapping and bias scores into the BMR could help correct for regions prone to variant-calling artefacts, while features like the number of palindrome loops and the fraction of mutations caused by AID per element could be used to adjust for locally-acting hypermutation processes.

When applied to the PCAWG data set, DriverPower called nearly twice as many non-coding driver events than coding ones, a ratio also observed by the PCAWG driver study. While this unbalanced ratio may reflect cancer biology, there is also the possibility that it reflects, at least in part, the technical challenge of sequencing and interpreting non-coding regions. Potential artefacts include systematic undercalling of somatic variants in non-coding regions^24^, a problem that could be rectified by deeper coverage. Another technical issue is raised by the fact that several non-coding candidates are only significant in the pan-cancer cohort, suggesting that the data set is statistically underpowered. To overcome this issue, we could either sequence more genomes or reduce the size of the set of test elements by narrowing it to functional motifs or conserved bases^57^. Lastly, functional impact score schemes are currently biased toward coding mutations; therefore, improved functional scoring schemes will also help us identify more functionally relevant non-coding cancer drivers in the future.

A comprehensive catalogue of coding and non-coding cancer drivers will accelerate the clinical translation of cancer genomic study to precision medicine. As more cancer genomes and more cancer types are sequenced, a general and accurate framework for computational driver discovery like DriverPower will become increasingly useful.

## Acknowledgements

This work was supported in part by the Government of Ontario.

## Methods

### Generation of cancer whole-genome somatic mutations

All DNA somatic single nucleotide variations (SNVs) and indels for 2,583 donors were obtained from the PCAWG project (somatic variant callset released October 2016)^9^. For our analysis, donors with hypermutated signatures were excluded (n=69, defined as more than 30 mutations per Mb). Otherwise, we used the same type-specific (n=29) and pan-cancer (all tumour samples except Skin-Melanoma, Lymph-NOS, Lymph-CLL and Lymph-BNHL) sample cohorts as the PCAWG Drivers and Functional Interpretation Group (PDFIG; **Supplementary Table 1**).

### Generation of simulated somatic mutations

We used three simulated datasets (Broad, DKFZ and Sanger simulations) from the PDFIG (described in detail at ref. xx). These simulations were made to capture the variation of background mutation rate and remove the signal of positive selection through permutations of observed somatic mutations.

### Generation of test and training genomic elements

We define a genomic element as the collection of genome coordinates that defines one specific functional region of interest. For example, the CDS element of *TP53* is the combination of all protein coding regions in *TP53*.

We used eight test element sets in our analysis, including the CDS (n=20,185), splice site (n=18,729), 5’UTR (n=19,369), 3’UTR (n=19,188), promoter (n=20,164), enhancer (n=30,816), lncRNA (n=5,580) and lncRNA promoter (n=5,373). All test element sets were obtained from the PCAWG project. GENCODE v19 was used as the reference gene model when building those sets^58^. Non-coding RNA annotations were collected from multiple sources as described.

We constructed genomic element training sets by randomly sampling genome coordinates from hg19, the build used for PCAWG. The length of each training element was sampled from the length distribution of test elements and multiplied by a factor of 3. Training elements overlapping test elements were removed. In total, 867,266 training elements were created and ˜54% (1,545,491,997 bp) of the genome was covered.

### Collection and generation of features

We collected 1,373 features in total. Details including data sources can be found at **Supplementary Table 2**. Nucleotide content features were calculated as the fraction of 2-mers and 3-mers in each genomic element. The number of 2-mers and 3-mers was counted directly from genome sequences (hg19). For raw features in bigwig format (typically genome wide signals), we calculated the average signal strength of covered bases in each element using the bigWigAverageOverBed (v2) utility from the UCSC genome browser^59^. For raw features in BED format (typically narrow peaks of ChIP-seq data), we calculated the percentage of bases intersecting BED for each element with the BEDTools (v2.24.0)^60^. All missing values in features were filled with 0.

### The DriverPower outline

The main steps of the DriverPower (v1.0.0) framework are summarized below. Details of each step are described in following sections. The difference between v1.0.0 and the version used in the PDFIG data analysis freeze (April 2017) is discussed in **Supplementary Note 2**.

1. Scale features and/or filter out excluded regions.
2. Build the background mutation rate model using the gradient boosting machine, or randomized lasso followed by binomial generalized linear models. The purpose of the BMR model is to estimate the expected number of mutations 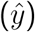 for any genomic element. Namely, we want to obtain 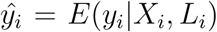 where *X*_*i*_ and *L*_*i*_ are the feature vector and length for the element *i*.
3. Conduct burden test with observed (*y*) and predicted 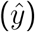 mutation counts, and perform multiple testing correction.
4. Adjust observed mutation counts (*y*) based on functional impact scores for nearly significant elements (q*<*0.25).
5. Re-assess the significance for nearly significant elements with functional adjusted mutation counts followed by multiple testing correction.

### Scaling of features

Features were scaled with RobustScaler from scikit-learn (version 0.18)^61^. Feature scaling was only conducted for randomized lasso and GLMs.

### Definition of excluded regions

In this study, all bases in the excluded regions were removed before any analysis. The excluded region consists of three sets: (1) all N bases and gaps in the hg19 genome (fetched from the UCSC table browser^12^); (2) ENCODE consensus excludable regions (the DAC Blacklisted Regions track and the Duke Excluded Regions track from the UCSC genome browser)^62^; (3) PCAWG low mappability regions (data retrieved from the PCAWG variant group). PCAWG low mappability regions are defined as regions callable in fewer than 556/1111 (˜50%) tumour-normal pairs. For each tumour-normal pair, a base is callable if there are more than 14/8 high quality reads in tumour/normal WGS. In total, 2,806,377,226 bp, or 96.41% of the genome are defined as callable.

### Feature selection with randomized lasso

To select features, we randomly sub-sampled 10% of the training set 500 times. Then for the *k*-th subset with size *N*_*k*_, the following model was fitted^63^:

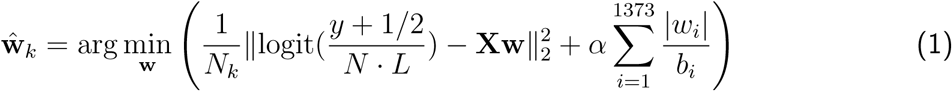

where *N* is the total number of donors in the dataset, **X** is the feature matrix, **w** is the weight vector, *α* is the regularization parameter, and *b*_*i*_ is the scaling factor. The regularization parameter *α* was determined by a 5-fold cross-validated lasso with 33% of the training data. For the *k*-th sub-sampling, the *i*-th feature was selected if 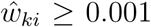. The final feature importance score was calculated as the fraction of times that a feature was selected. Only features with score *>* 0.5 were used in the GLM BMR model.

### Prediction of the BMR with GLM

When using the generalized linear model, we modelled the observed mutations in each genomic element with a binomial distribution, that is

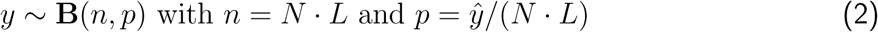

where *y* is the observed mutation count and 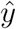 is the estimated mutation count. We used the binomial generalized linear model to obtain 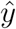 with the logit link function, that is

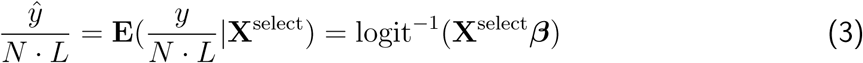

where **X**^select^ is the selected feature matrix and *β* is the regression coefficient vector.

### Prediction of the BMR with GBM

We trained the gradient boosting machine with XGBoost^64^. All features were used in model training. The negative Poisson log-likelihood was chosen as the objective function and ln(*N · L*) of elements were used as offset (*i.e.*, `base_margin` in XGBoost). Other non-default parameters used in DriverPower were as follows: `eta=0.05, max depth=8, subsample=0.6, max delta step=1.2, early stop rounds=5 and nrounds=5000`. The feature importance for GBM is measured by the improvement in accuracy brought by a feature across all trees. XGBoost returns feature importance that sums up to 1 for all features. We also normalized the importance to a [0,1] scale (*i.e.*, importance relative to the most important feature).

### Evaluation of two BMR models

We evaluated both models with 1Mb autosome bins (n=2,521) and training genomic elements (n=867,266) defined above. The 1Mb elements have been used in many studies^14,65,15^. For both elements, we obtain the predicted mutation rate by 5-fold cross validation (CV). For 1Mb elements, we used 4-fold data for model training and 1-fold data for model evaluation. For training elements, we use 1-fold data to train the model and the rest to evaluate. As per previous work, we used *R*^2^ score and Pearson’s *r* as evaluation metrics^15^. Standard error of the mean (SEM) for *R*^2^ and *r* was calculated from 5-fold CV.

### Calculation of element functional impact scores

Four different functional scores were used in this analysis^16,17,18,19^. For CDS, CADD (SNVs and indels, v1.3), DANN (SNVs) and EIGEN (SNVs) scores were used. CADD indel scores were generated with the CADD web interface for all observed indels in the PCAWG dataset. For splice site, CADD and DANN scores were used. For non-coding elements, the CADD, DANN and LINSIGHT (SNVs and indels) score were used. Then the following steps were used to calculate the functional impact score per genomic element. Firstly, raw scores were retrieved for all observed mutations in the dataset. Secondly, all raw scores were converted to phred-like scores by −10 log_10_(rank/*N*_*m*_), where *N*_*m*_ is the number of observed mutations having scores. Thirdly, for each genomic element, its functional score *S* was calculated as:

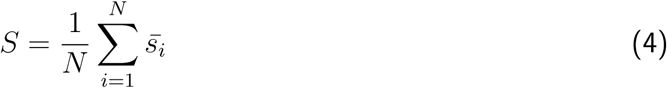

where *N* is the number of donors and 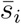 is the average functional impact score for the *i*th donor.

### Adjustment of the mutation count

To compensate for the unbalanced number of mutations among samples, instead of using the mutation count per element directly we used the geometric mean of mutation count and sample count. That is, we use the balanced count 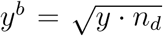 instead of *y* directly for significance test, where *n*_*d*_ is the number of mutated donors. Based on the motivation that not all mutations should be weighted the same, the balanced mutation count *y*^*b*^ was then adjusted for nearly significant elements (raw q-value < 0.25) by a functional weight *w*, that is *y*^*f*^ = *w ⋅ y*^*b*^, where *y*^*f*^ is the functionally adjusted mutation count. For the element *j*, the functional weight *w*_*j*_ was calculated based on its functional score *S*_*j*_ and a threshold score *S*_*T*_:

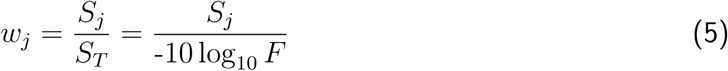

The threshold score *S*_*T*_ is controlled by a single parameter *F* between 0 and 1, and can be interpreted as the fraction of functionally relevant variants among all observed variants. Parameter tuning of *F* can be found at **Supplementary Note 1**.

### Assessment of the element significance

For each element, we calculated 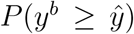 as the raw p-value and *P* 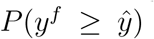 as the function-adapted p-value. Since over-dispersion has been documented in burden based methods and can affect the driver discovery accuracy^22^, here we performed a regression-based overdispersion test for each tumour cohort using the training set^66^. Based on the result of the overdispersion test, we calculated the raw and function-adpated p-values by following a binomial distribution or a negative binomial distribution:

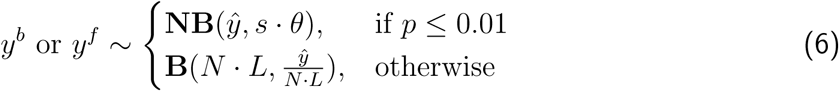

where *p* and *θ* are the p-value and dispersion parameter estimated from the overdispersion test, and *s* is the scaling factor for *θ* used to accommodate the discrepancy between test and training set in terms of the dispersion level. We used *s* = 3 for lymphomas and *s* = 1 otherwise in this analysis.

### Multiple testing correction

In all cases, q-values were generated by the Benjamini–Hochberg procedure^67^. We chose q<0.1 as the significant level and q<0.25 as the nearly-significant level. For each element set, multiple testing correction was performed for each tumour cohort (cohort q-value) and across all tumour cohorts (global q-value). Cohort q-values were used in functional adjustment and global q-values were used to define the final driver list.

### Generation of reference cancer drivers

Reference cancer drivers were used to benchmark the performance of DriverPower. Three reference sets were used: (1) the COSMIC Cancer Gene Census (v82, n=567); (2) the PCAWG consensus driver candidates (PCAWG-consensus; n=157 for coding and n=26 for non-coding); (3) the PCAWG raw integrated driver candidates (PCAWG-raw; n=193 for coding and n=79 for non-coding). PCAWG-consensus (q-value post-filtering < 0.1) is a set of highly confident non-coding drivers and subjected to multiple stringent filters as described. PCAWG-raw (q-value pre-filtering < 0.1) is a superset of PCAWG-consensus and includes non-coding drivers that were not subjected to the filtering process. PCAWG-raw driver candidates that are mutated in fewer than three samples were removed in this analysis. For promoter and 5’UTR candidates in the PCAWG consensus drivers, we reversed the filtering for overlapping elements (*i.e.*, one element is selected over the overlapping element based on prior knowledge). For example, we kept both the promoter and the overlapping 5’UTR of *WDR74* in this analysis; in the PCAWG consensus set, the *WDR74* promoter is preferentially selected over its 5’UTR.

### Benchmarking of DriverPower

We compared coding and non-coding driver candidates called by DriverPower to driver candidates called by six other published driver detection tools (ActiveDriverWGS^25^, ExInAtor^20^, LARVA^22^, MutSig^24^, ncdDetect^21^ and oncodriveFML^23^). Driver calls for 26 single tumour cohorts (no Skin-Melanoma, Lymph-CLL and Lymph-BNHL) were retrieved from the PCAWG driver group. For each method, we removed driver candidates that are mutated in fewer than three samples. We used precision (*TP/(TP+FP)*), recall (*TP/(TP+FN)*) and F1-score (`2*Precision*Recall/(Precision+Recall)`) as performance metrics.

For CDS, we used the CGC gene set as the gold standard. For each method, true positive genes were defined as genes presented in the gold standard set and the precision was then calculated as the fraction of true positive genes among all called genes. For recall, since we can’t accurately know the expected set of driver genes that should be called for each tumour cohort in the dataset, a lower-bound approximation was used instead. The lower-bound approximation was estimated by taking the union of all true positive genes identified by each method and the recall was then calculated as the fraction of true positive genes called among the lower-bound approximation.

For gene splice sites, the same gold standard gene set and benchmark method as CDS were used. Due to data availability, the comparison was only performed for ncdDetect, oncodriveFML and DriverPower.

For promoters, enhancers, 3’UTRs and 5’UTRs, because the number of non-coding driver candidates is small, four element sets were benchmarked together. No data for ExInAtor is available for this comparison. For each tumour cohort, true positive driver elements were defined as elements called by at least three methods. The calculation of precision, recall and F1-score was then identical as for the CDS and splice site.

### Somatic copy number and structural variations analysis

We used SCNA (including GISTIC2.0 results) and SV call sets released January 2017^68^. The copy number status (loss, neutral or gain) of a region is classified based on the difference between the absolute copy number of the region and the genome-wide ploidy of the donor. For gene-level structural variations, we calculated the number of breakpoints per gene (including CDS, splice sites, UTRs and promoters) per donor.

### Differential expression analysis

We used the upper quartile normalized gene expression (FPKM-UQ) released May 2016^69^. When comparing the expression difference between two groups of donors, we fitted the following quasi-Poisson family generalized linear model and then employed the likelihood ratio test to obtain p-values for mutational status: FPKM-UQ ˜ MUT + SCNA + [Tissue], where MUT is the mutational status (0 for unmutated donors and 1 for mutated donors), SCNA is the somatic copy number status (-1, 0 or 1 for copy number loss, neutral or gain, respectively) and Tissue is the tumour tissue type. The tissue type was only used for pan-cancer comparison for the adjustment of tumour types.

### WES data analysis

We obtained two whole exome sequencing datasets through the Genomic Data Common (GDC)^70^: TCGA-PAAD (35,321 somatic mutations across 180 samples) and TCGA-LIHA (56,208 somatic mutations across 364 samples). We chose public MuTect2 variants from GDC. Variant coordinates were lifted from hg38 to hg19 with the UCSC liftOver tool. Only CADD scores were used to detect drivers. For TCGA-PAAD, GBM models trained from Panc-AdenoCA of the PCAWG data were used. For TCGA-LIHA, GBM models trained from Liver-HCC were used.

### Code availability

The source code for DriverPower (written mainly in Python 3) is available at https://github.com/smshuai/DriverPower. It is distributed under GNU General Public License 3.0, which allows for reuse and redistribution.

### Data availability

The PCAWG data of simple somatic mutations, coding and non-coding cancer driver calls, RNA expression values, copy number somatic mutations and structural somatic mutations are available at the ICGC DCC (https://dcc.icgc.org/pcawg).

## Supplementary Information

### Supplementary Notes

#### Supplementary Note 1 Parameter tuning

As noted in the **Method** section, a single parameter *F* (the functional score threshold) was used to control the degree of functional adjustment in DriverPower. Functional scores used in DriverPower were positively correlated with the functional impact of mutations within elements. The parameter *F* controls the threshold score (*S*_*T*_ = -10 log_10_ *F*) and must be within the interval (0,1]. The parameter *F* is negatively correlated with *S*_*T*_. Genomic element with score *S > S*_*T*_ will be up-weighted; while element with score *S < S*_*T*_ will be down-weighted. Since DriverPower uses phred-scale scores, *F* can be interpreted as the proportion of functionally relevant variants among all observed variants. For instance, *F* = 0.01 means *S*_*T*_ = 20 so elements with functional score *S* > 20 (i.e., rank top 1% in phred-like scale) will receive a functional weight *>*1 and gain additional significance from functional impact information. Using larger *F* will result in smaller *S*_*T*_, and more elements will obtain additional significance, we hence expect to obtain more driver candidates. Empirically speaking, larger *F* will cause higher recall but lower precision as illustrated in **Supplementary Fig. 16**.

The choice of *F* is dependent on the score scheme in use and/or the element set and tumour cohort in test. Here to avoid overfitting, we divided all 2583 donors into training donor set (N=1,117) and test donor set (N=1,136) to choose *F* for four score schemes (CADD, DANN, EIGEN and LINSIGHT) and three element types (CDS, splice site and other non-coding sites) separately. For CDS and splice site, we used the COSMIC cancer gene census (CGC) as the gold standard set for parameter tuning. For other non-coding sites, we used the PCAWG-raw as the gold standard set for parameter tuning. Precisions were calculated as the fraction of hits in the gold standard set and pseudo-recalls were calculated as the number of significant candidates with functional adjustment in gold standard over the number of nearly-significant candidates without functional adjustment in gold standard. This calculation of pseudo-recall measured the relative performance of DriverPower to its personal best. We observed that optimal parameters for the training donor set also worked in a similar way for the test donor set. Parameters learnt here were used for driver discovery in this analysis.

#### Supplementary Note 2 Difference from the PCAWG freeze

The list of coding and non-coding drivers produced by this analysis (v1.0.0) differs slightly from the DriverPower results included in the PDFIG data analysis freeze due to the following reasons:

1. Hypermutated samples were not removed for the PCAWG freeze.
2. Only the binomial test was used in the PCAWG freeze.
3. The BMR model used in the PCAWG freeze was randomized lasso followed by GLM and in this analysis was gradient boosting machines.
4. Only CADD and EIGEN scores are used in the PCAWG freeze, and element p-value is the minimal p-value generated from two scores.

The threshold scores used for CADD and EIGEN in the PCAWG freeze were cohort-specific. Namely, 85%-95% percentile score of each tumour cohort was used as threshold.

Around 83% of candidates identified by v1.0.0 were also in PCAWG freeze (**Supplementary Fig. 17a,c**). For coding driver discovery, the current version has higher precision and lower recall (**Supplementary Fig. 17b**). For non-coding driver discovery, four candidates are only significant in the current version, including the promoter of lncRNA *RMRP* in Breast-AdenoCA and Stomach-AdenoCA, as well as *LEPROTL1* promoter in Bladder-TCC and *TERT* promoter in CNS-Oligo (**Supplementary Fig. 17d**). In addition, the removal of hypermutated samples and the incorporation of negative binomial test alleviated the inflation issue in melanomas and lymphomas.

### Supplementary Tables

Supplementary Table 1 Summary of tumour cohorts

Supplementary Table 2 Summary of genomic features

Supplementary Table 3 Coding and non-coding driver candidates called by DriverPower

Supplementary Table 4 DriverPower-exclusive driver candidates

### Supplementary Figures

**Supplementary Figure 1 Heterogeneity at cohort and donor levels**. The violin plot shows the number of somatic mutations (log-scale) per donor across all tumour types. Hypermutated donors are included in the figure. Circles in the violin plot represent median values.

**Supplementary Figure 2 Heterogeneity at the locus level.** The average number of mutations per cohort per Mb are shown for chromosome 22. Black lines show the mean mutation rate per 1Mb interval. Filled ribbons show the mutation rate within 1 standard deviation.

**Supplementary Figure 3 Features correlation matrix.** The heatmap shows the Spearman’s rank correlation coefficient matrix between 1,373 features. Rows and columns are ordered by feature subgroups (40 subgroups in total). Barplot at right of the heatmap shows the Spearman’s rho between features and mutation rates in training data for pan-cancer.

**Supplementary Figure 4 Expected versus GLM predicted mutation rate.** Mutation rates are the number of non-coding mutations within non-overlap 1Mb genome windows and normalized to per Mb per donor. Predictions are made from the randomized lasso and the generalized linear model. The *R*^2^ (coefficient of determination) and *r* (Pearson’s correlation coefficients) are calculated with 5-fold cross validation. Note that only selected features are used in the GLM.

**Supplementary Figure 5 Expected versus GBM predicted mutation rate.** Mutation rates are the number of non-coding mutations within non-overlap 1Mb genome windows and normalized to per Mb per donor. Predictions are made from the gradient boosting machines. The *R*^2^ (coefficient of determination) and *r* (Pearson’s correlation coefficients) are calculated with 5-fold cross validation. All features are used in the GBM.

**Supplementary Figure 6 Model comparison.** (**a**) Variance explained for all tumour cohorts in 1Mb genome elements. (**b**) Variance explained for all tumour cohorts in the training element set. Error bars show the standard error of the mean (SEM). SEM is calculated from 5-fold cross validation. GBM is the gradient boosting machine. GLM is the generalized linear models.

**Supplementary Figure 7 Feature importance rank.** All features are divided into subgroups (x-axis). Maximum importance of each group per tumour cohort (including pan-cancer) is represented as a point. Violin plot shows the distribution and median feature importance per subgroup among tumour cohorts. (**a**) Feature importance measures by the randomized lasso. Red line shows the 0.5 cutoff used in the generalized linear model. (**b**) Feature importance measures by the GBM (log-scale).

**Supplementary Figure 8 Top features associated with cell lines having similar origins.** Barplot shows the importance of 15 replication timing features for CNS-GBM, Liver-HCC and Breast-AdenoCA. X-axis shows the cell line name where the repli-seq experiment is conducted (ENCODE). Red line shows the 0.5 cutoff used in the feature selection. Feature weights from the GBM are normalized to [0, 1].

**Supplementary Figure 9 Elemental functional impact scores.** (**a**) Functional impact scores across all sets of elements for four different score schemes (CADD, DANN, EIGEN and LIN-SIGHT). Random elements are the training elements used in the model. (**b**) Spearman’s correlation matrix for each set of elements (pairwise complete).

**Supplementary Figure 10 P-value quantile-quantile plots.** The p-value QQ-plots are shown for all test element types across three simulated datasets generated by the PDFIG and the observed PCAWG dataset. For the observed dataset, both raw p-values (blue) and function-adapted p-values (red) are shown. For the Broad, DKFZ and Sanger simulations, only raw p-values are shown and elements with q-value<0.1 are labelled. Each QQ-plot contains p-values from all tumour cohorts as the FDR control is performed in this way. Elements without mutations are removed in the plot. For better visualization, -log10 observed p-values are capped at 16.

**Supplementary Figure 11 Functional adjustment improves CDS driver discovery.** Black horizontal and vertical dashed lines show the q-value = 0.1 cutoffs. Only significant genes are labelled with colors corresponding to reference gene sets. Function-adapted q-values are q-values with the usage of functional impact scores. Genes in the top-right quadrant are significant in both raw and functional adjusted results, in the bottom-left quadrant are significant in neither. Genes in the top-left are significant only after functional adjustment and genes in the bottom-right are removed after functional adjustment.

**Supplementary Figure 12 Comparison of DriverPower and MutSig for CDS results.** The Venn diagram shows the relationship of true positive CDS results for DriverPower and Mut-Sig. Significant genes present within the COSMIC CGC are considered to be true positive calls. DriverPower identifies 21 and MutSig identified 23 true positive genes exclusively.

**Supplementary Figure 13 Benchmark result of splice sites.** The left panel shows the precision and recall for each method according to the results on 26 tumour cohorts. The right panel shows a heatmap of significant candidates identified by each method. Significant candidates in CGC are filled with blue and not in CGC are filled with black.

**Supplementary Figure 14 DriverPower-exclusive driver candidates overview.** (**a,b**) CDS of *EEF1A2* in Eso-AdenoCA. (a) is a mutation lollipop plot of *EEF1A2* in Eso-AdenoCA. (b) shows the somatic copy number status for *EEF1A2* in Eso-AdenoCA or non-Eso-AdenoCA (other) samples; the p-value is from the Fisher’s exact test. (**c**) A lollipop plot for CDS of *MEF2B* in Lymph-BNHL. (**d**) A violin plot for differential expression between Lymph-BNHL samples with mutated (MUT) *MEF2B* and non-mutated (WT) *MEF2B.* (**e**) Lollipop plots for CDS and splice site of *SGK1* in Lymph-BNHL. (**f**) Lollipop plots for *GPR126* enhancer in Breast-AdenoCA and Bladder-TCC. Lollipop plots show the distribution and classes of mutations (legends on top). In the lollipop plot, element is shown as a rectangular box and blocks of a disjoint element are separated by vertical dashed lines. Text within the box shows the name, type and length (bp) of elements. Arrow below the element box indicates the direction of element.

**Supplementary Figure 15 WES driver discovery result.** P-value Q-Q plots for two TCGA whole-exome sequencing datasets. Only significant genes are labelled (q<0.1). Blue points and labels indicate candidates within reference driver sets; red points and labels indicate likely false positive hits.

**Supplementary Figure 16 Parameterization of DriverPower.** Each panel shows a parameter (*F*) search for one of the score schemes (CADD, DANN, EIGEN and LINSIGHT) and one of the element sets (CDS, splice site and other non-coding elements). The best *F* is indicated below each panel title. For each search path, the left end point is the smallest *F* searched and the right end point is the largest *F* searched; points correspond to *F* searched and triangles correspond to the best *F*. Search paths are colored for training (blue) and test (red) donor sets (see **Supplementary Note 1** for more details).

**Supplementary Figure 17 Comparison of PCAWG freeze and v1.0.0. (a,b)** Comparison of coding driver candidates for non-melanoma/lymphoma tumours. (a) is the venn plot showing the relationship between CDS results from PCAWG freeze and CDS results from this analysis (v1.0.0). (b) shows the number and fraction of candidates called for two versions. Columns in (b) are colored by reference gene sets. **(c,d)** Comparison of non-coding driver candidates for non-melanoma/lymphoma tumours. (c) is the venn plot showing the relationship of non-coding results (3’UTR, 5’UTR, promoter, enhancer, lncRNA and lncRNA promoter) between PCAWG freeze and v1.0.0. (d) shows all non-coding candidates that are unique to PCAWG freeze (n=7) or v1.0.0 (n=4). See **Supplementary Note 2** for more details.

